# Contact of mobile *cis*-Golgi network with ER exit sites executes COPI-mediated receptor recycling

**DOI:** 10.64898/2026.10.06.756745

**Authors:** Ramon Rodriguez-Prieto, Yasuyuki Suda, Takuro Tojima, Susana Sabido-Bozo, Kazuo Kurokawa, Akihiko Nakano, Manuel Muñiz

## Abstract

COPI-mediated retrograde transport is essential for maintaining homeostasis between the endoplasmic reticulum (ER) and the Golgi in eukaryotic cells. However, its spatiotemporal dynamics remain unclear. In budding yeast, ER export occurs through a “hug-and-kiss” mechanism, wherein mobile *cis*-Golgi network (CGN) elements transiently approach ER exit sites (ERES) to capture cargo before departing to deliver it to the Golgi. Here, using high-resolution 3D and 4D live imaging, we show that the COPI-mediated recycling of a cargo receptor from the CGN back to the ER occurs during the hug-and-kiss event. Furthermore, this recycling is complete before the CGN detaches from the ERES, suggesting that COPI-mediated retrograde transport is spatiotemporally coupled to ER export during a single and transient CGN-ERES association. Consistent with this model, when COPI function is impaired, the receptor fails to return efficiently to the ER and instead remains in the CGN after separation from the ERES. Additionally, COPI impairment prolonged CGN-ERES association, suggesting that COPI function also facilitates CGN detachment from ERES. Thus, we propose that rapid COPI-mediated recycling drives CGN maturation directly at the ERES interface, enabling the CGN to detach and proceed along the secretory pathway.

## Introduction

The eukaryotic secretory pathway begins with the transport of newly synthesized proteins from the endoplasmic reticulum (ER) to the *cis*-Golgi network (CGN), an intermediate compartment localized between the ER and Golgi (Hsu, Yuan et al. 1991). Historically, this conserved structure has also received several other names, such as the ER-Golgi intermediate compartment (ERGIC/IC) in mammals (Schweizer et al. 1990, Saraste and Svenson 1991) and the Golgi entry core compartment (GECCO) in plants (Schweizer, Fransen et al. 1990, Ito, Uemura et al. 2018). To unify the divergent terminology, we recently proposed adopting CGN as the standard term (Nakano 2026). From the CGN, proteins are transferred to the Golgi stack for post-translational modification and sorting to their final destinations (Bonifacino and Glick 2004, Szul and Sztul 2011, Barlowe and Helenius 2016, Gomez-Navarro and Miller 2016, Farhan, Raote et al. 2025). The conventional model posits that the initial ER-to-CGN traffic relies on transport carriers formed by the assembly of the COPII protein coat at specialized ER exit sites (ERES) (Kurokawa and Nakano 2019, Melero, Boulanger et al. 2022, Downes, Flood et al. 2026). This anterograde membrane flux is counterbalanced by a COPI vesicle-mediated retrograde pathway. This retrograde transport is essential because it recycles membranes, the machinery required for subsequent rounds of anterograde transport and retrieves escaped ER-resident or misfolded proteins (Letourneur, Gaynor et al. 1994, Spang 2013, Arakel, Huranova et al. 2019, Downes, Flood et al. 2026, Yang, Yang et al. 2026). Consequently, maintaining homeostasis within the early secretory pathway should require a strictly regulated balance between COPII-mediated anterograde and COPI-dependent retrograde traffic. Despite its fundamental importance to cellular physiology, whether and how these opposing membrane fluxes are coordinated in space and time remains poorly understood.

Advances in high-spatiotemporal-resolution live-cell microscopy have revealed a novel mechanism for ER-to-CGN transport (Nakano 2013, Kurokawa, Okamoto et al. 2014). Unlike the conventional model, where free COPII vesicles travel through the cytosol toward the CGN, super-resolution confocal live imaging microscopy (SCLIM) has shown that anterograde transport in the budding yeast *Saccharomyces cerevisiae* is driven by a highly mobile CGN (Kurokawa, Okamoto et al. 2014, Tojima, Suda et al. 2024). This mobile compartment transiently approaches the ERES to directly capture cargo, presumably by fusing to COPII carriers. Following cargo loading, the CGN detaches and moves away, completing the anterograde transport step. While this dynamic hug-and-kiss mechanism ensures efficient and safe cargo delivery, it raises the critical question of how and when the COPI retrograde transport operates to recycle the anterograde machinery back to the ER.

In this study, we addressed this question by using SCLIM to analyze the dynamics of a cargo receptor shuttling between the ER and the CGN. Cargo receptors are transmembrane proteins that escort specific cargo out of the ER during COPII-mediated transport (Dancourt and Barlowe 2010, Barlowe and Helenius 2016). Upon reaching the CGN, the cargo must dissociate to continue along the secretory pathway, while the empty receptor returns to the ER via COPI-dependent retrograde transport. This continuous recycling makes these receptors an ideal model system for studying retrograde transport within the early secretory pathway. We find that COPI-mediated receptor recycling occurs during the transient association between the CGN and the ERES in the hug-and-kiss event. Furthermore, we observed that the receptor completes its full recycling before the CGN detaches from the ERES, suggesting that COPI-mediated retrograde transport is spatiotemporally coupled to ER export during a single, transient CGN-ERES association. Moreover, we also reveal that COPI facilitates the detachment of the CGN from ERES. Our findings suggest that COPI-driven receptor recycling initiates CGN maturation upon ERES association and promotes its subsequent detachment to progress along the secretory pathway.

## Results

### The p24 cargo receptor localizes at steady state at ERES and ERES-associated CGN

To analyze the dynamic behavior of cargo receptors cycling between the ER and the CGN, we first assessed the steady-state localization of the anterograde cargo receptor p24 complex in living yeast cells using high-resolution 3D imaging. The p24 complex is a heteromeric protein complex composed of members of the conserved p24 protein family (Marzioch, Henthorn et al. 1999, Strating and Martens 2009, Hirata, Nihei et al. 2013, Pastor-Cantizano, Montesinos et al. 2016, Tashima, Hirata et al. 2022). These abundant type I transmembrane proteins feature cytosolic COPII and COPI coat-binding signals that facilitate their bidirectional trafficking between the ER and the CGN (Sohn, Orci et al. 1996, Rojo, Pepperkok et al. 1997, Füllekrug, Suganuma et al. 1999, Belden and Barlowe 2001, Aguilera-Romero, Kaminska et al. 2008). The p24 complex has been shown to play a role as selective anterograde cargo receptor by promoting the efficient ER export of specific cell surface proteins (Muñiz, Nuoffer et al. 2000, Bonnon, Wendeler et al. 2010, Castillon, Aguilera-Romero et al. 2011, Fujita, Watanabe et al. 2011, Manzano-Lopez, Perez-Linero et al. 2015). We performed simultaneous three-color, 3D SCLIM visualization of the p24 complex subunit Emp24 tagged with GFP (D’Arcangelo, Crissman et al. 2015), together with ERES and CGN markers. As an ERES marker, we used the outer COPII coat subunit Sec13 tagged with iRFP, given that the ERES is the site of COPII coat assembly (Kurokawa, Okamoto et al. 2014, Kurokawa and Nakano 2019). As an CGN marker, we utilized the syntaxin Qa-SNARE Sed5 tagged with mCherry, which predominantly localizes to the CGN (Tojima, Suda et al. 2024). Both markers displayed their characteristic distributions in *S. cerevisiae* cells: Sec13-iRFP-marked ERES appeared as numerous scattered puncta throughout the nuclear and peripheral ER, while the CGN exhibited a dispersed distribution of Sed5-positive cisternae in the cytoplasm (Figure 1A) (Tojima, Suda et al. 2024). While the majority of Emp24-GFP colocalized with or was resided immediately adjacent to ERES, some fraction was also observed to colocalize with the CGN (Figure 1B). Because the CGN can be found either associated with ERES during hug-and-kiss events or physically separated from them before and after the event, we sought to determine which of these two CGN subpopulations contained Emp24-GFP. We first confirmed that approximately 55% of CGN structures were intimately associated with ERES, consistent with the previously described hug-and-kiss mechanism for cargo capture (Kurokawa, Okamoto et al. 2014, Tojima, Suda et al. 2024). Interestingly, Emp24-GFP preferentially colocalized with these ERES-associated CGNs (80%), likely representing the pool of Emp24-GFP transferred from ERES to the CGN via anterograde transport during the hug-and-kiss interaction (Figures 1D and 1E). Panel 1 of Figure 1F shows a typical example of ERES-associated CGN containing Emp24-GFP. Conversely, Emp24-GFP was present at a significantly lower abundance in free, physically separated cisternae (20%), as shown in Figures 1D and 1E. Panel 2 of Figure 1F depicts a representative example of a free, ERES-separated CGN lacking Emp24-GFP. Collectively, these findings suggest that Emp24-GFP is captured by the CGN during the transient CGN-ERES association and is rapidly recycled back to the ER via retrograde transport before the CGN departs.

**Figure 1:**
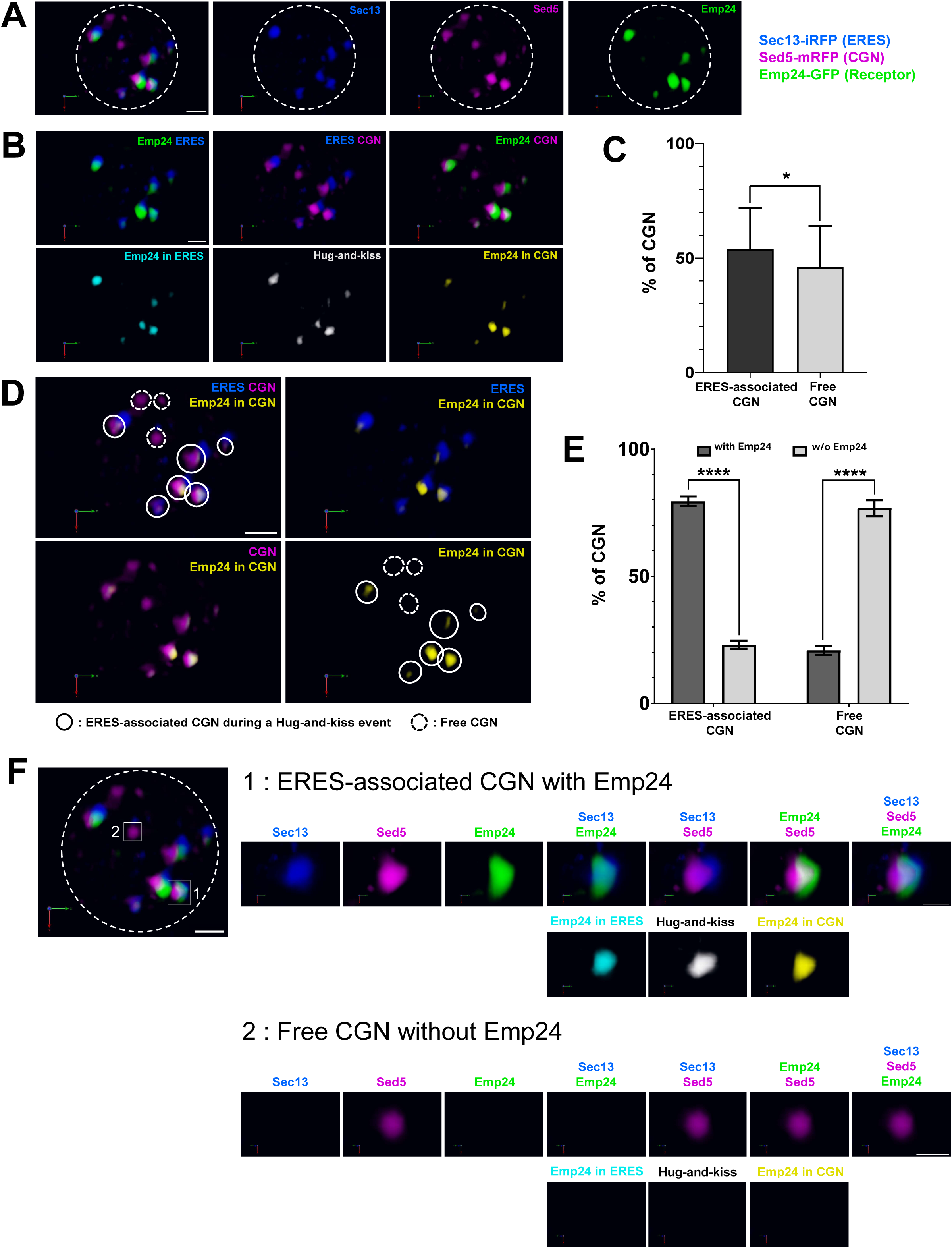
The cargo receptor Emp24 localizes at steady state to ERES and transiently ERES-associated CGN. Wild-type cells expressing Sec13-iRFP (ERES, blue), Sed5-mCherry (CGN, magenta), and Emp24-GFP (cargo receptor, green) were observed at 24°C by SCLIM. **(A)** Representative 3D images of merged markers (ERES, CGN, and cargo receptor markers, left) and each individual marker (ERES, CGN and cargo receptor) are shown. Scale bar, 1 µm. **(B) Top panels** show 3D merged images of Emp24 and ERES, ERES and CGN, and Emp24 and CGN markers. **Lower panels** show colocalization channels to display only the Emp24-GFP present in the ERES (cyan), the ERES-associated CGN (hug-and-kiss event, white) and the Emp24-GFP present in CGN (yellow). Scale bar, 1 µm. **(C)** Quantification of micrographs described in (B) showing colocalization between ERES and CGN markers. Average percentage per cell of ERES-associated CGN (hug-and-kiss events) and free CGN physically separated from ERES. n = 792 in 63 cells in three independent experiments. Error bars = SD. Two-tailed, unpaired t test. *P < 0.05. **(D) Right upper panel:** 3D merged images of ERES (blue), CGN (magenta) and Emp24-GFP present in CGN (yellow). **Left upper panel:** 3D merged images of ERES (blue) and Emp24-GFP present in CGN (yellow). **Right lower panel:** 3D merged images of Emp24-GFP present in CGN (yellow). **Left lower panel**: Emp24 in ERES-associated CGN (orange). Open arrowheads: ERES-associated CGN. Gray filled arrowheads: Free CGN physically separated from ERES. Scale bar, 1 µm. **(E)** Quantification of micrographs described in (D) showing colocalization. Average percentage of ERES-associated CGN (hug-and-kiss events) and free CGN physically separated from ERES with or without Emp24-GFP. n = 792 in 63 cells in three independent experiments. Error bars = SD. 2-way ANOVA test. ***P < 0.0001. **(F) Left panel** shows the same representative 3D image of merged ERES, CGN and cargo receptor markers displayed in (A) with two selected individual CGN denoted (CGN 1: ERES-associated CGN containing Emp24-GFP and CGN 2: free ERES-separated CGN without Emp24-GFP). Scale bar, 1 µm. **Right panels** show enlarged 3D images of selected CGN 1 and CGN 2. Individual and merged markers (ERES, CGN, and cargo receptor) and colocalization images are shown. Scale bar, 0,5 µm.

### The p24 cargo receptor returns to the ER prior to the CGN detaching from ERES

Our 3D SCLIM data showed the absence of Emp24-GFP in the free CGN, suggesting that this cargo receptor is retrieved back to the ER before the CGN separates from the ERES (Figure 1). To test this hypothesis directly, we monitored CGN detachment events by using simultaneous three-color, 4D (3D time-lapse) SCLIM visualization of Emp24-GFP together with the ERES marker Sec13-iRFP and the CGN marker Sed5-mCherry. In 4D imaging of transient hug-and-kiss interactions, Sec13-iRFP and Sed5-mCherry initially colocalized (t = 0 s). As the CGN subsequently detached and moved away from the ERES, this colocalization was lost (t = 20 s; Figure 2A). Notably, colocalization analysis also revealed that the portion of Emp24-GFP initially present in the ERES-associated CGN (t = 0 s) did not remain as the compartment detached (t = 20 s; Figure 2A). The line scan analysis shown in Figure 2B confirmed that the CGN harbors Emp24-GFP while associated with the ERES, but loses it upon dissociation, implying that Emp24 must be returned to the ER during the hug-and-kiss event. Statistical analysis of colocalization over time across 214 independent detachment events clearly showed that Emp24-GFP is recycled before the CGN separates from the ERES (Figure 2C). Altogether, these results support that cargo receptors enter the CGN and return to the ERES, completing a full cycle of anterograde and retrograde transport, during the hug-and-kiss interaction.

**Figure 2:**
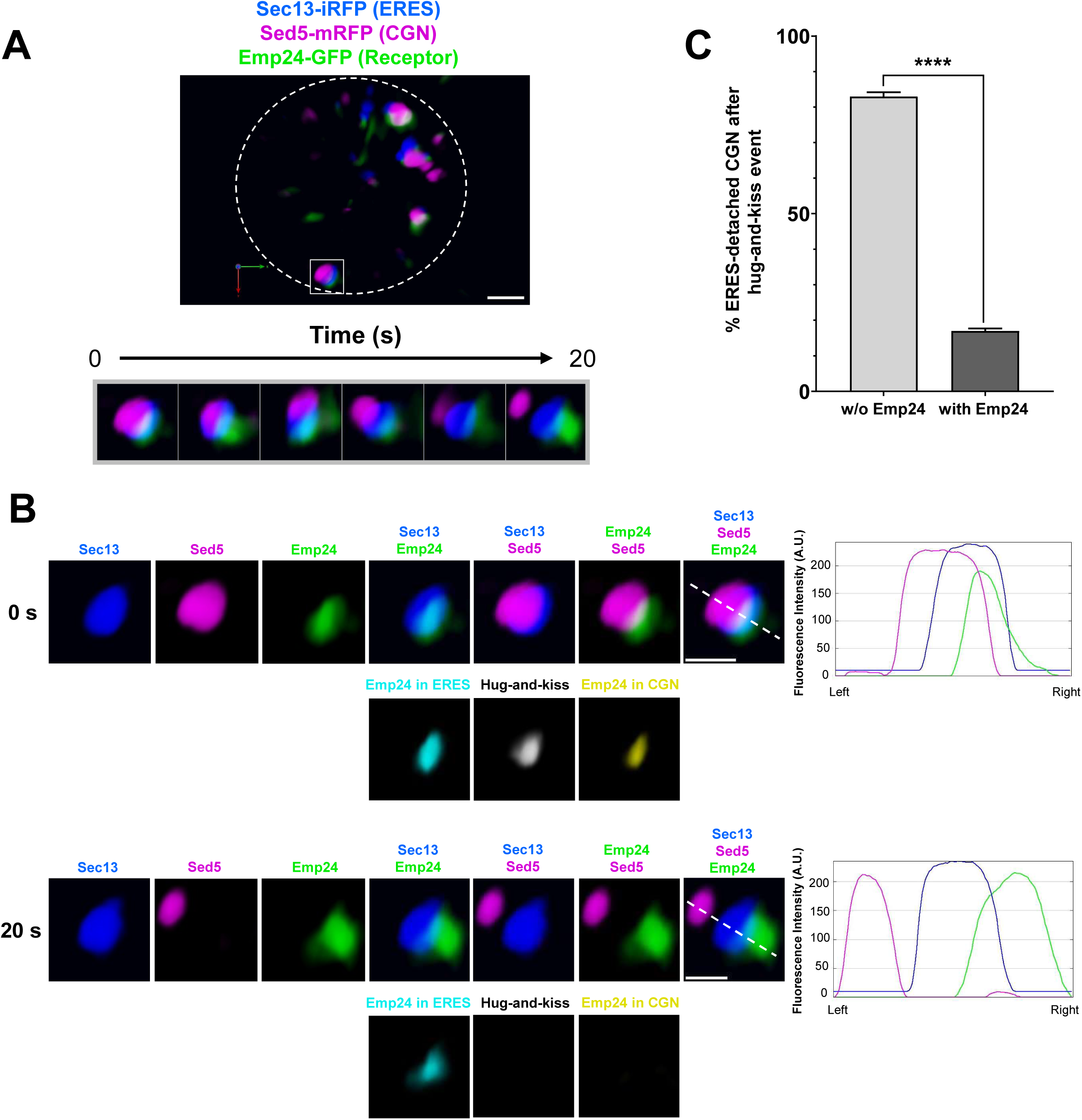
Retrograde transport of Emp24-GFP from the CGN to the ER occurs before CGN detaches from ERES. Three-color 4D observation of CGN separation from ERES in living yeast cells. Wild-Type cells expressing Sec13-iRFP (ERES, blue), Sed5-mCherry (CGN, magenta), and Emp24-GFP (cargo receptor, green) were observed at 24°C by SCLIM. **(A) Top panel:** A representative 3D image of merged ERES, CGN, and cargo receptor markers is shown. Scale bar, 1 µm. **Middle panels** show 4D (3D time-lapse) images of a selected CGN (boxed area in the top panel). 3D images were reconstructed from 21 optical slices 0,2 µm apart around the center of the cell taken at 4-s intervals. Merged images of ERES, CGN, and cargo receptor markers are shown. **Lower panels** show selected 4D (3D time-lapse) images (boxed images in the middle panel) corresponding to 0 s and 20 s time frame points. Individual and merged markers (ERES, CGN, and cargo receptor) and colocalization images are shown. Scale bar, 0,5 µm. Emp24 is present in the ERES-associated CGN (t = 0 s) but absent from the ERES-detached CGN (t = 20 s). **(B)** Quantification of micrographs from (A) showing colocalization between Emp24-GFP and the CGN marker during CGN detachment from ERES. The graph displays the average percentage of Emp24-GFP-positive, ERES-associated CGN structures that subsequently detach from ERES, either losing or retaining the Emp24-GFP signal. n = 214 in 33 cells in three independent experiments. Error bars = SD. Two-tailed, unpaired t test. ***P < 0.0001.

### In COPI-impaired cells, the p24 cargo receptor remains within the detached CGN upon its dissociation from ERES

The COPI coat drives the retrograde transport pathway that recycles back cargo receptors from CGN to the ER. Therefore, to further verify that cargo receptor recycling occurs during the hug-and-kiss event and before the CGN detaches from ERES, we examined using SCLIM the p24 cargo receptor behavior in the COPI mutant *ret1-1*. The temperature-sensitive *ret1-1* mutant has a lesion in the α subunit of coatomer and even at permissive temperature (24°C) shows already a defect in the CGN-to-ER retrograde transport (Letourneur, Gaynor et al. 1994). We reasoned that if Emp24-GFP is recycled exclusively from ERES-associated CGN, then Emp24-GFP in *ret1-1* strain should remain trapped in the CGN after its detachment from the ERES. We tested this prediction by performing both simultaneous three-color, 3D and 4D SCLIM visualization of Emp24-GFP, Sec13-iRFP and Sed5-mCherry in *ret1-1* cells at 24°C. The 3D SCLIM analysis first confirmed that approximately 45% of the CGN population is associated with ERES, while 55% remains free and separated from ERES in the *ret1-1* cells (Figure 3C). Notably, the majority of Emp24-GFP (60%) was present in this free CGN pool, in contrast to the wild-type strain, where only about 20% of Emp24-GFP resided in free CGN (Figure 3E). Panels of Figure 3F show a representative example of a free, ERES-detached CGN containing Emp24-GFP in the *ret1-1* strain.

**Figure 3:**
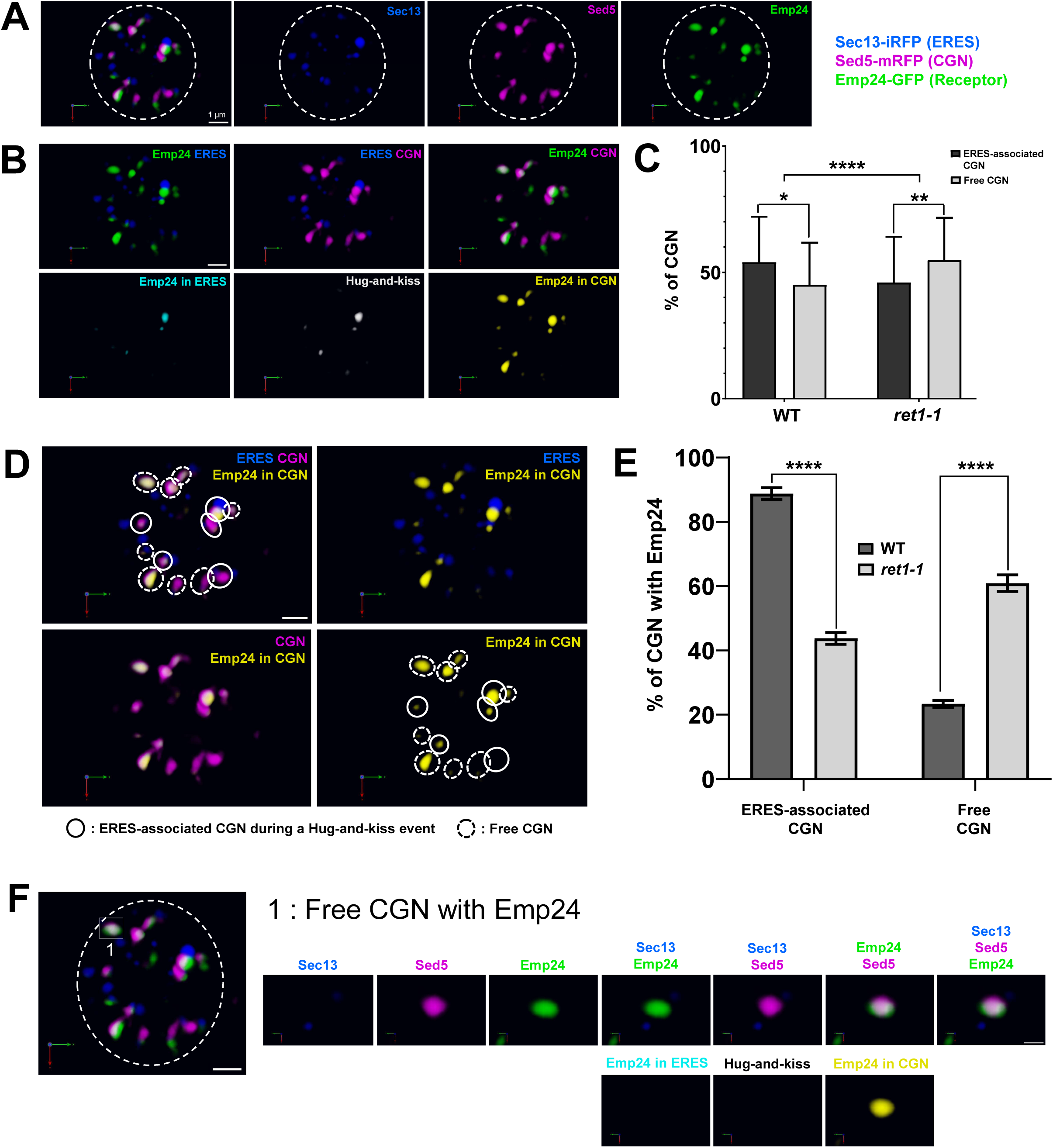
The presence of Emp24-GFP at the free CGN increases at steady state in COPI-deficient cells. COPI thermosensitive mutant (*ret1-1*) cells expressing Sec13-iRFP (ERES, blue), Sed5-mCherry (CGN, magenta), and Emp24-GFP (cargo receptor, green) were observed at 24°C by SCLIM. **(A)** Representative 3D images of merged markers (ERES, CGN, and cargo receptor markers, left) and each individual marker (ERES, CGN and cargo receptor) are shown. Scale bar, 1 µm. **(B) Top panels** show 3D merged images of Emp24 and ERES, ERES and CGN, and Emp24 and CGN markers. **Lower panels** show colocalization channels to display only the Emp24-GFP present in the ERES (cyan), the ERES-associated CGN (hug-and-kiss event, white) and the Emp24-GFP present in CGN (yellow). Scale bar, 1 µm. **(C)** Quantification of micrographs described in (B) showing colocalization between ERES and CGN markers. Average percentage per cell of ERES-associated CGN (hug-and-kiss events) and free CGN physically separated from ERES in wild-type (WT) and *ret1-1* cells. n = 963 in 71 cells in three independent experiments. Error bars = SD. Two-way ANOVA test with Tukey’s multiple comparisons test; ****P < 0.0001 for interaction, *P < 0.05, **P < 0.01. **(D) Left upper panel:** 3D merged images of ERES (blue), CGN (magenta) and Emp24-GFP present in CGN (yellow). **Right upper panel:** 3D merged images of ERES (blue) and Emp24-GFP present in CGN (yellow). **Left lower panel:** 3D merged images of Emp24-GFP present in CGN (yellow). **Right lower panel:** Emp24 in ERES-associated CGN (orange). Open arrowheads: ERES-associated CGN. Gray filled arrowheads: Free CGN physically separated from ERES. Scale bar, 1 µm. **(E)** Quantification of micrographs described in (D) showing colocalization. Average percentage of ERES-associated CGN (hug-and-kiss events) and free CGN physically separated from ERES with or without Emp24-GFP in wild-type (WT) and *ret1-1* cells. N = 963 in 71 cells in three independent experiments. Error bars = SD. 2-way ANOVA test. ***P < 0.0001. **(F) Left panel** shows the same representative 3D image of merged ERES, CGN and cargo receptor markers displayed in (A) with a selected individual CGN denoted (boxed area) representing an example of an ERES-detached CGN containing Emp24-GFP Scale bar, 1 µm. **Right panels** show enlarged 3D images of selected CGN. Individual and merged markers (ERES, CGN, and cargo receptor) and colocalization images are shown. Scale bar, 0,5 µm.

Next, we conducted 4D SCLIM analysis to monitor the p24 cargo receptor dynamics upon CGN detachment from ERES in *ret1-1* cells. 4D SCLIM tracked the physical dissociation of the CGN from the ERES following hug-and-kiss events, marked by the loss of initial Sec13-iRFP and Sed5-mCherry colocalization. Subsequent analysis showed that in the *ret1-1* cells the Emp24-GFP fraction initially present in the ERES-associated CGN (t = 0 s) remained trapped within the compartment upon detachment (t = 20 s; Figure 4A). Line-scan profiles (Figure 4B) confirmed this retention in CGN upon ERES separation, highlighting a failure of Emp24 retrograde trafficking to the ER during these transient interactions in *ret1-1* cells. Statistical quantification across 236 detachment events further proved that, in contrast to the wild-type strain, Emp24-GFP fails to recycle and instead escapes with the departing CGN in *ret1-1* mutant cells (Figure 4C). These results confirm that COPI-mediated receptor recycling occurs before the CGN dissociates from ERES. Since recycling by COPI has been shown to drive cisternal maturation (Ishii, Suda et al. 2016, Kurokawa, Osakada et al. 2019), our data reveal that the CGN undergoes maturation upon ERES association.

**Figure 4:**
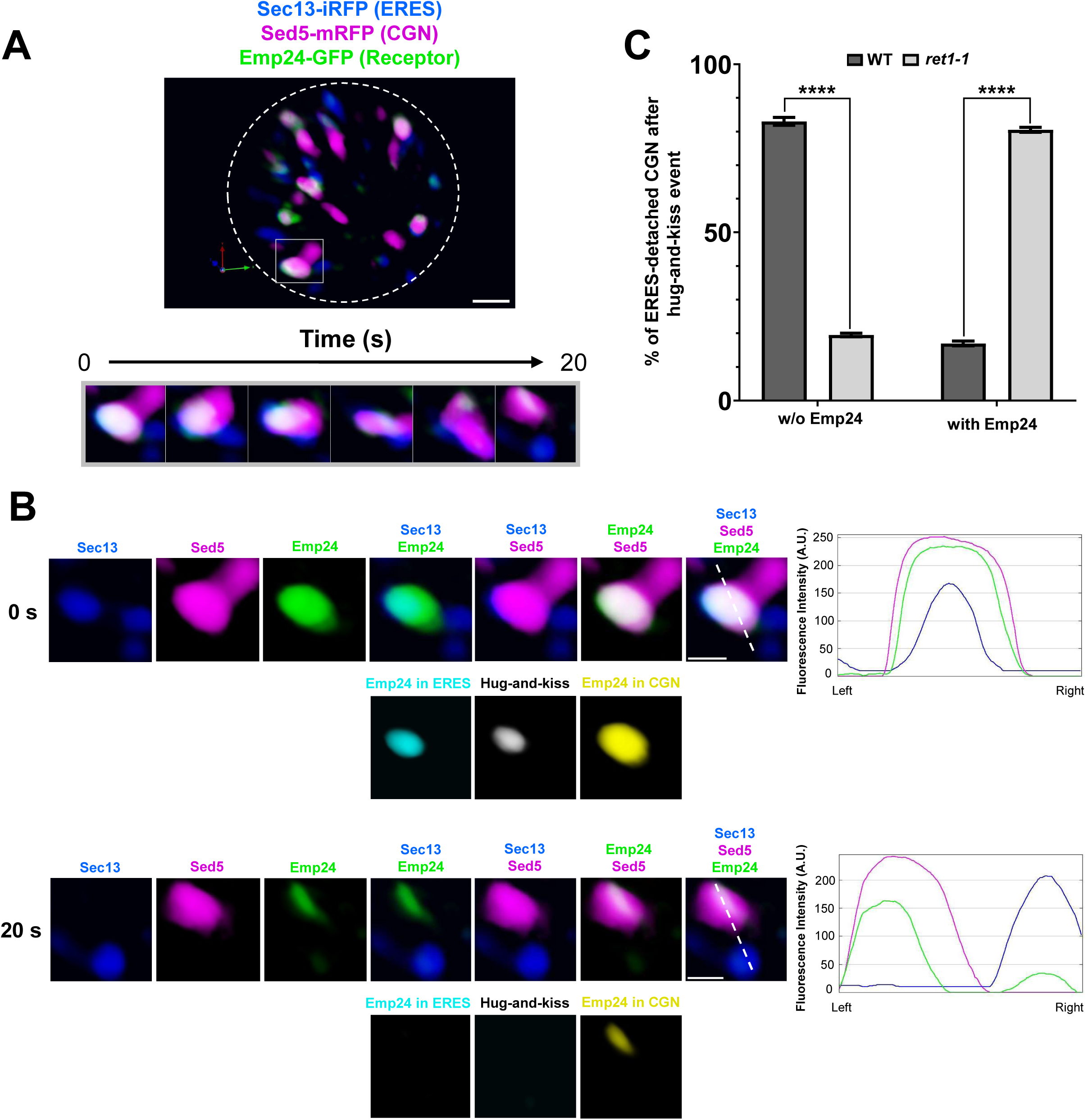
Emp24-GFP fails to efficiently return to the ER and remains in the CGN following detachment from the ERES in COPI-deficient cells. Three-color 4D observation of CGN separation from ERES in living yeast cells. COPI thermosensitive mutant (*ret1-1*) cells expressing Sec13-iRFP (ERES, blue), Sed5-mCherry (CGN, magenta), and Emp24-GFP (cargo receptor, green) were observed at 24°C by SCLIM. **(A) Top panel:** A representative 3D image of merged ERES, CGN, and cargo receptor markers is shown. Scale bar, 1 µm. Middle panels show 4D (3D time-lapse) images of a selected CGN (boxed area in the top panel). 3D images were reconstructed from 21 optical slices 0,2 µm apart around the center of the cell taken at 4 s intervals. Merged images of ERES, CGN, and cargo receptor markers are shown. **Lower panels** show selected 4D (3D time-lapse) images (boxed images in the middle panel) corresponding to 0 s and 20 s time frame points. Individual and merged markers (ERES, CGN, and cargo receptor) and colocalization images are shown. Scale bar, 0,5 µm. Emp24 is present in the ERES-associated CGN (t = 0 s) but absent from the ERES-detached CGN (t = 20 s). **(B)** Quantification of micrographs from (A) showing colocalization between Emp24-GFP and the CGN marker during CGN detachment from ERES. The graph displays the average percentage of Emp24-GFP-positive, ERES-associated CGN structures that subsequently detach from ERES, either retaining or losing the Emp24-GFP signal in wild-type (WT) and *ret1-1* cells. n = 236 in 39 cells in three independent experiments. Error bars = SD. 2-way ANOVA test. ***P < 0.0001.

### COPI-driven maturation facilitates CGN detachment from ERES

During our SCLIM 4D analysis of Emp24-GFP dynamics in the *ret1-1* mutant, we noticed an apparent prolonged association between the CGN and ERES. To investigate this further, we quantified and compared the CGN-ERES dissociation kinetics in both wild-type and *ret1-1* strains. As shown in Figure 5, wild-type cells predominantly exhibited transient CGN-ERES associations that peaked at 4 seconds and declined progressively. Conversely, *ret1-1* mutants displayed a marked reduction in these short-term associations (4–12 s). This decrease was accompanied by a concomitant increase in prolonged associations (>32 s) and the emergence of non-dissociating compartments. This finding indicates that proper COPI function is required for efficient CGN detachment from the ERES. We conclude that, after capturing cargo via a hug-and-kiss interaction, the CGN compartment must undergo COPI-driven cargo receptor recycling to efficiently separate from the ERES and proceed maturation along the secretory pathway.

**Figure 5:**
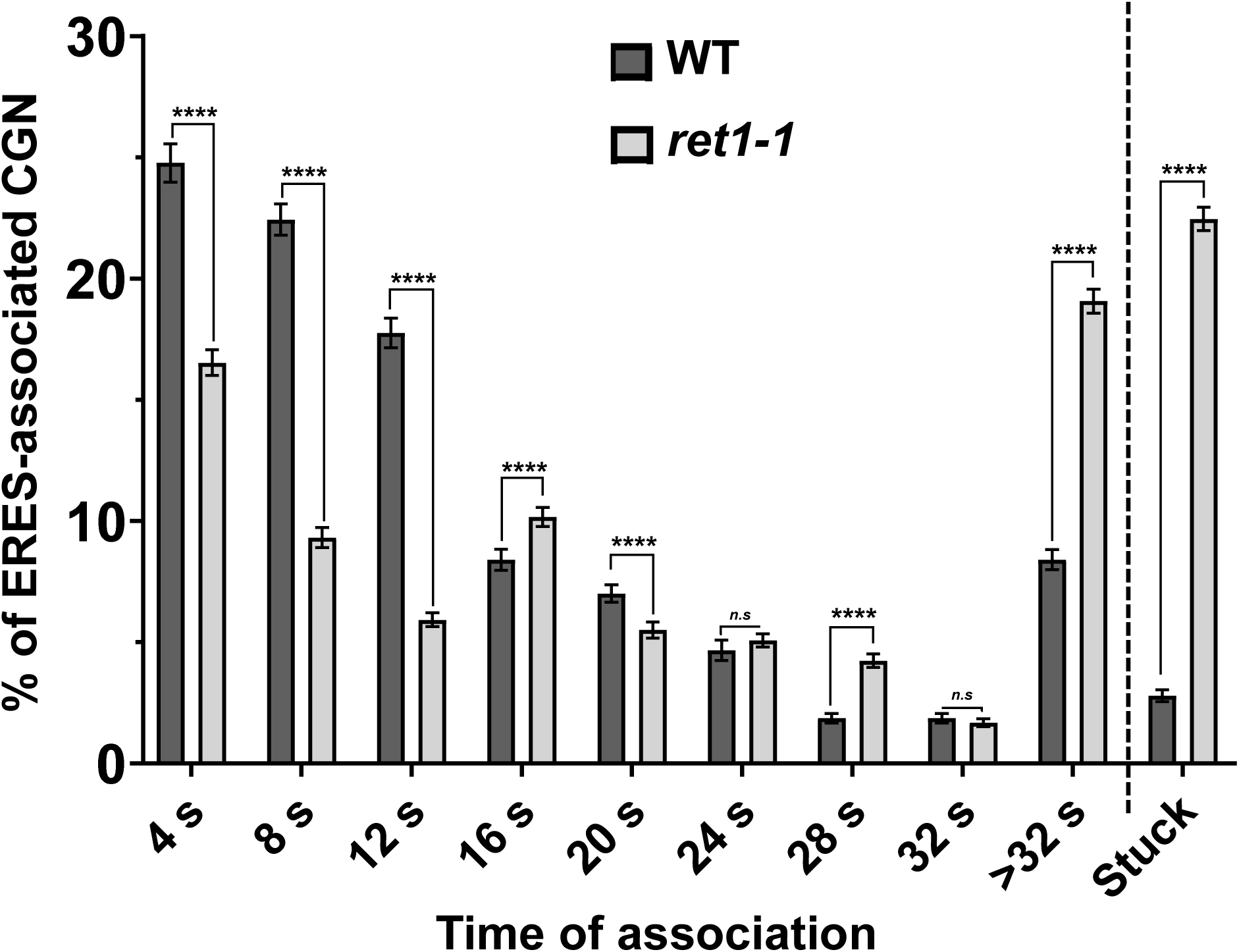
Persistent association between CGN and ERES in COPI-deficient cells. Quantification of the temporal dynamics of CGN-ERES association. The graph displays the percentage of ERES-associated CGNs as a function of association duration in wild-type (WT, dark gray) and *ret1-1* mutant (light gray) cells. WT cells exhibit predominantly transient interactions that peak at 4 s and progressively decline over time. In contrast, the *ret1-1* mutant displays a reduction in short-term associations (4-12 s) along with a concomitant increase in prolonged interactions (>32 s) and stable, non-dissociating compartments (“Stuck”). N = 214 in 36 WT cells and n = 236 in 39 *ret1-1* cells in three independent experiments. Error bars = SD. 2-way ANOVA test. ****P < 0.0001.

## Discussion

COPI-mediated retrograde transport is essential for cellular homeostasis, yet its spatiotemporal dynamics within the early secretory pathway remain unclear (Farhan, Raote et al. 2025). To investigate this fundamental question, we utilized SCLIM, a powerful high-spatiotemporal-resolution imaging technology (Kurokawa, Ishii et al. 2013). We previously employed SCLIM to reveal the hug-and-kiss cargo ER export mechanism in the budding yeast *S. cerevisiae*, wherein mobile CGN elements transiently associate with ERES to capture secretory cargo and then leave away (Kurokawa, Okamoto et al. 2014, Tojima, Suda et al. 2024). In this study, we used SCLIM to monitor the retrieval dynamics of the p24 cargo receptor from the CGN to the ER. The 3D SCLIM analysis revealed that, at steady state, Emp24-GFP preferentially localizes to ERES and ERES-associated CGN in wild-type cells, but accumulates in free, ERES-separated CGN in a COPI-deficient mutant. Moreover, 4D SCLIM dynamics demonstrated that in wild-type cells, Emp24-GFP returns to the ER during the transient CGN-ERES association phase, before detachment. In COPI-deficient cells, however, this retrograde transport is abrogated, leaving Emp24-GFP stranded in the CGN following its departure from the ERES. These results show that COPI-mediated retrograde transport of cargo receptors from the CGN to the ER is executed and spatiotemporally coupled to ER export during the transient association between the CGN and ERES (Figure 6). Given that cargo receptors such as the p24 receptor have been shown to facilitate vesicle budding by binding the COPI coat through specific sequence motifs (Bremser, Nickel et al. 1999, Aguilera-Romero, Kaminska et al. 2008, Mitrovic, Ben-Tekaya et al. 2008), we suggest that, upon releasing their secretory cargo, receptors selectively recruit COPI and spatially segregate into a distinct CGN zone. This spatial segregation of receptors would facilitate the COPI vesicle budding required for their rapid return to the contiguous ER membrane during the hug-and-kiss event.

**Figure 6:**
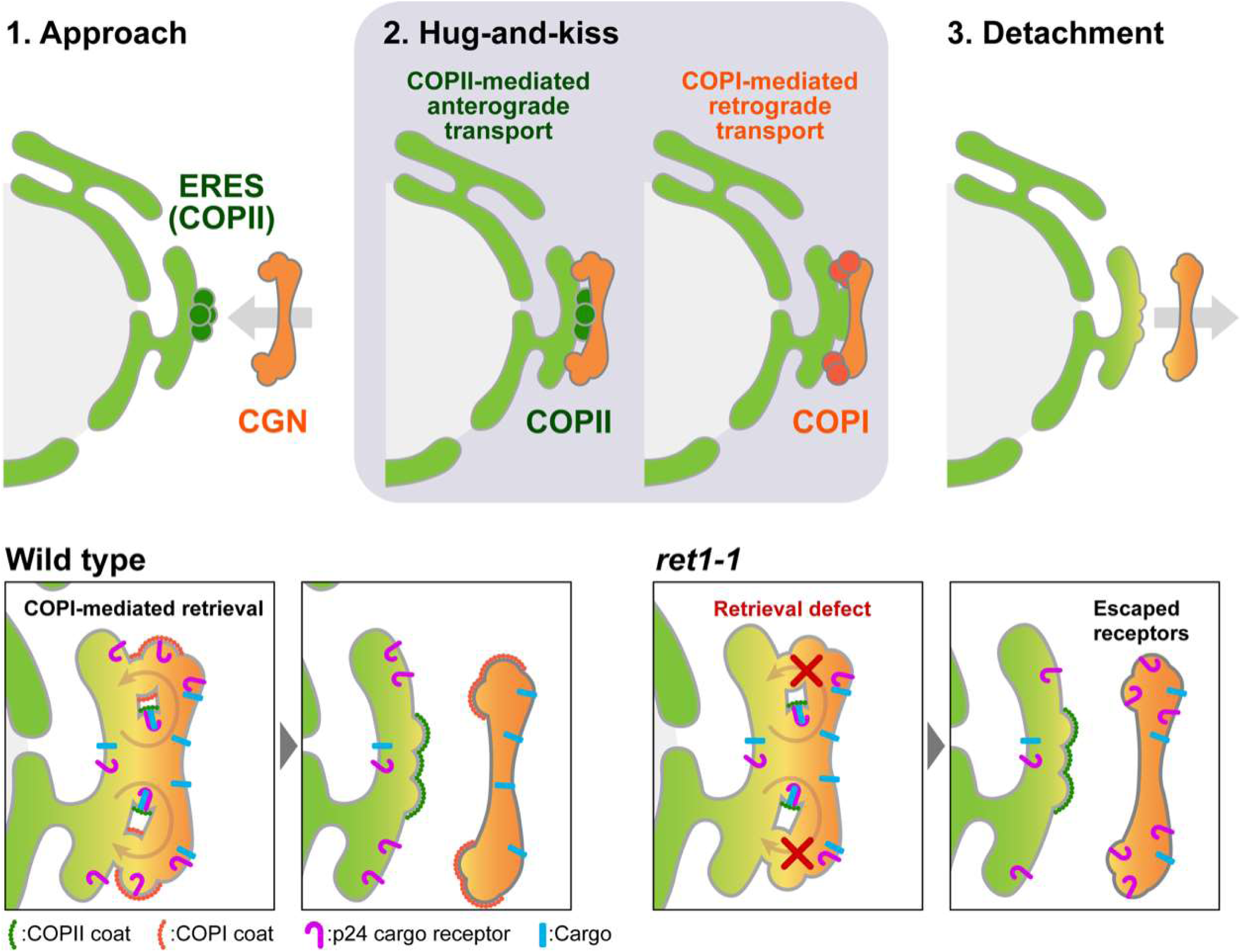
A model for coordinated bidirectional transport of a cargo receptor between ER and CGN by a hug-and-kiss action. First, the mobile CGN approaches and associates with the ERES to capture cargo bound to its transmembrane receptor. Next, while the CGN remains associated with the ERES, the receptor segregates from the cargo and returns empty to the ER via the COPI-mediated retrograde pathway. Once this receptor retrieval is complete, the cargo-bearing CGN detaches from the ERES to progress along the secretory pathway through cisternal maturation. The SCLIM resolution used in this study cannot distinguish whether COPII-mediated export and COPI-dependent recycling occur via lipid vesicles, via buds that fuse to form transient tunnels, or a combination of both. However, both mechanisms remain compatible with our model.

Our previous studies and those of others have shown that in the budding yeast the secretory pathway downstream of the CGN operates via cisternal maturation (Losev, Reinke et al. 2006, Matsuura-Tokita, Takeuchi et al. 2006, Ishii, Suda et al. 2016, Kurokawa, Osakada et al. 2019, Tojima, Suda et al. 2024). In this mechanism, rather than moving between static compartments, secretory cargo remains within a single membrane-bound compartment that undergoes biochemical transformation. Specifically, the CGN sequentially matures into a *cis*-Golgi cisterna, a *trans*-Golgi cisterna, and finally the *trans*-Golgi network (TGN), facilitating complete post-translational cargo processing. This progressive shift in cisternal identity is driven by COPI-mediated retrograde transport (Papanikou, Day et al. 2015, Ishii, Suda et al. 2016). By continually recycling resident proteins, such as early Golgi glycosylation enzymes, back to the preceding, younger cisterna, the compartment matures around the stationary cargo, propelling it forward through the secretory pathway. Remarkably, our results suggest that the first step of this maturation pathway occurs while the CGN remains associated to the ERES, during the hug-and-kiss event. Upon receptor-cargo dissociation, the CGN immediately initiates COPI-mediated retrograde transport to recycle cargo receptors back to the ER. This rapid recycling enables the receptors to initiate new rounds of ER export, while the cargo stays trapped within the maturing CGN. Thus, our findings position the yeast CGN as a dynamic sorting or recycling compartment that initiates COPI-driven maturation at the ERES interface, detaches, and progressively matures into distinct Golgi compartments.

Furthermore, beyond this role in receptor recycling during the hug-and-kiss event, our findings also reveal that COPI facilitates CGN detachment from the ERES. This is supported by the delayed detachment kinetics observed in the *ret1-1* mutant. Such a delay likely impairs cargo progression through the secretory pathway, providing a mechanistic explanation for the defective anterograde ER-to-Golgi transport characteristic of COPI mutants (Novick, Field et al. 1980, Gaynor and Emr 1997, Sütterlin, Doering et al. 1997). Although the precise mechanism remains to be fully elucidated, this reduced detachment aligns with our previous observations in budding yeast, where the mobility of Sed5-labeled compartments was restricted in *ret1-1* cells, further reinforcing the role of COPI in regulating CGN dynamics (Ishii, Suda et al. 2016). Furthermore, this mobile CGN that has dissociated from the ERES could be viewed as an “anterograde” carrier, somewhat analogous to the mobile COPI structures recently observed in mammalian cells that eventually separate from the ERES (Weigel, Chang et al. 2021).

This study further emphasizes that close spatial proximity between the ER and CGN, achieved by the hug-and-kiss mechanism, is a prerequisite for coordinated bidirectional trafficking between these compartments. Strongly supporting this notion, recent studies in yeast and mammalian cells have shown that ER acceptor sites (ERAS), where COPI retrograde carriers undergo tethering and fusion, are intimately associated with ERES (Schröter, Beckmann et al. 2016, Roy Chowdhury, Bhattacharjee et al. 2020). Nevertheless, our finding that COPI-driven receptor recycling relies on the CGN-ERES association raises two key questions. The first concerns the physical nature of the COPI retrograde transport intermediates involved. One possibility is that discrete, COPI-coated lipid vesicles bud from the CGN and traverse the narrow spatial gap to fuse with ERAS, where the short distance ensures rapid and efficient delivery. Alternatively, the tip of a COPI bud or a tubule formed at the CGN could transiently fuse with the ERAS membrane, creating a continuous connection until subsequent fission at the CGN separates them. This fission perhaps might contribute to the CGN detachment from the ER. Such a “kiss-and-run” mechanism could even coexist with standard vesicular transport, depending on the dynamic distance between the CGN and ERES/ERAS (Melero, Boulanger et al. 2022, Farhan, Raote et al. 2025, Downes, Flood et al. 2026, Mironov and Beznoussenko 2026). Because the spatial resolution of our SCLIM analysis cannot currently discriminate between these distinct mechanisms, further studies will be required to resolve these possibilities.

The second fundamental question concerns the molecular mechanism orchestrating the hug-and-kiss event. Uncovering this will require re-examining transport machinery components from a new perspective. Indeed, our observations suggest that, like the COPI coat, the long coiled-coil tethering factor Uso1 promotes CGN detachment from ERES, given that the *uso1-1* mutant displays a similarly prolonged compartment association (Kurokawa, Okamoto et al. 2014). Additionally, condensates driven by intrinsically disordered regions (IDRs) may act as a “molecular glue” to stabilize the ERES-CGN association and limit carrier diffusion (Ruiz, Bogus et al. 2026). In mammalian cells, for example, the IDR-enriched protein TFG forms condensates that restrict COPII carrier mobility (Bogus, Wegeng et al. 2025). Although budding yeast lacks a direct TFG ortholog, other IDR-enriched proteins like the golgin Grh1 could serve a similar function (Rebane, Ziltener et al. 2020, G Oliveira, Silva et al. 2026). Future studies must clarify how the transient CGN-ERES contact spatiotemporally coordinates COPI-mediated retrograde transport with ER export, and whether this mechanism is evolutionarily conserved in plants and mammals.

## Methods

### Yeast strains, plasmids, and culture conditions

Strains of *S. cerevisiae* used for this work are MMY1828 (MAT**a** ADH1p-2xmRFP-SED5(SnaBI)::TRP1 SEC13-iRFP::natNT2 ade2-101_ochre ura3-52 lys2-801_amber trp1-Δ63 his3-Δ200 leu2-Δ1) and MMY1829 (*MAT*α *ret1-1 ADH1p-2xmRFP-SED5(SnaBI)::TRP1 SEC13-iRFP::natNT2 ade2-101_ochre ura3-52 trp1-Δ63 his3-Δ200 leu2-Δ1*) strains used for live-cell imaging by SCLIM were constructed in the YPH499 background (Brachmann et al., 1998). The insertion of *SEC13-iRFP::natNT2* and *ADH1p-2xmRFP-SED5(SnaBI)::TRP1* into the yeast genome was performed by PCR-mediated gene replacement (Janke et al., 2004; Longtine et al., 1998) and verified by PCR and fluorescence microscopy. *EMP24-sfGFP* was expressed under the control of its own promoter on the *CEN* plasmid YCplac111 (pJD062, a gift from E. Miller (D’Arcangelo, Crissman et al. 2015)). For microscopic observation, the yeast cells were grown in selective medium (0.67% yeast nitrogen base without amino acids and 2% glucose) with appropriate supplements. The cells were harvested at an early-to-mid logarithmic phase and then seeded onto glass coverslips coated with concanavalin A.

### Live-cell imaging by SCLIM

Yeast cells were grown in MCD medium (0.67% yeast nitrogen base without amino acids [Difco Laboratories], 0.5% casamino acids [Difco Laboratories], and 2% glucose) with appropriate supplements. For live imaging, cells were grown to a mid-log phase at 24°C. Cells were immobilized on glass slides using concanavalin A and imaged by SCLIM. SCLIM was developed by combining Olympus model IX-71 inverted fluorescence microscope with a UPlanSApo 100× NA 1.4 oil objective lens (Olympus), a high-speed and high signal-to-noise ratio spinning-disk confocal scanner (Yokogawa Electric), a custom-made spectroscopic unit, image intensifiers (Hamamatsu Photonics) equipped with a custom-made cooling system, magnification lens system for giving 266.7× final magnification, and three EM-CCD cameras (Hamamatsu Photonics) for green (sfGFP), red (2xmRFP), and infrared (iRFP) observation (Kurokawa et al., 2013). Image acquisition was executed by custom-made software (Yokogawa Electric). For 3D and 4D live imaging, we collected optical sections spaced 100 or 200 nm apart in stacks by oscillating the objective lens vertically with a custom-made piezo actuator. Z stack images were converted to 3D voxel data and processed by deconvolution with Volocity (Perkin Elmer) using the theoretical point-spread function for spinning-disk confocal microscopy. Colocalization analysis was performed using Costes’ method thresholding (Costes et al., 2004) of Volocity software. Line-scan analysis was performed using Fiji (ImageJ version 1.54p, National Institutes of Health, Bethesda, MD, USA; Schindelin et al., 2012).

## Statistical analysis

Statistical significance was determined using GraphPad Prism software. For the two-tailed Student’s t test and ordinary two-way analysis of variance (ANOVA) test, differences among groups were considered significant for P < 0.05 (*).

## Acknowledgments

This manuscript is dedicated to the memory of our dear colleague and friend, Kazuo Kurokawa, who passed away during the preparation of this work. We thank Miho Waga, Kalai Madhi Muniandy, Sofia Rodriguez-Gallardo, and Antonio Cordones-Romero for their technical assistance.

## Author contributions

K. Kurokawa, Y. Suda performed the experiments; R. Rodriguez-Prieto, Y. Suda and S. Sabido-Bozo analyzed the data; K. Kurokawa and S. Sabido-Bozo made the strains; M. Muñiz wrote the manuscript; R. Rodriguez-Prieto, Y. Suda, T. Tojima and A. Nakano revised the manuscript; K. Kurokawa, M. Muñiz and Y. Suda designed the experiments and supervised the project.

## Abbreviations

3D: three-dimensional
4D: four-dimensional
ER: endoplasmic reticulum
ERES: ER exit sites
COPII: coat protein complex II
CGN: cis-Golgi Network
COPI: coat protein complex I
ERGIC: ER-Golgi intermediate compartment
IC: Intermediate Compartment
GECCO: Golgi entry core compartment
TGN: trans-Golgi network
SCLIM: super-resolution confocal live imaging microscopy

